# scRetinaDB: A Comprehensive Database of Single-Cell and Spatial Omics from Cross-Species Retinas

**DOI:** 10.1101/2025.10.11.681556

**Authors:** Miaoxiu Tang, Fan Jiang, Andrei-Florian Stoica, JingJing Chen, Jiongliang Wang, Kejun Yao, Zhaoming Chen, Xueli Xu, Jie Wang

## Abstract

The retina is essential for encoding visual signals, and its dysregulation can lead to retinal diseases. Recent advances in single-cell and spatial sequencing technologies have yielded extensive omics data from retinal tissues across species and biological conditions. However, existing retinal omics data are dispersed across various repositories without uniform processing, which limits integrative analysis. To address this we developed scRetinaDB (https://casapp.dnayun.com/scretina/), a comprehensive resource that aggregates single-cell RNA sequencing (scRNA-seq), single-cell assay for transposase-accessible chromatin using sequencing (scATAC-seq), and spatial RNA sequencing (spRNA-seq) data from retinas across species and diverse biological conditions. The database comprises over 2.79 million retinal cells collected from 453 scRNA-seq datasets spanning 34 studies, 17 species and 27 biological conditions. For each species, these scRNA-seq datasets were integrated to construct a retinal cell atlas. In addition, scRetinaDB also contains 107 scATAC-seq and 18 spRNA-seq datasets from human and mouse retinas. The scRetinaDB website provides four major modules separately for browsing species-specific omics data, searching cross-species omics profiles, performing analyses of cell type annotation and cell similarity analysis, and downloading preprocessed multi-omics datasets. Overall, scRetinaDB is a valuable resource for retinal single-cell and spatial omics, advancing cross-species studies particularly in vision research.

## Background

The retina is a critical part of the eye that performs phototransduction, processes visual signals, and transmits them to the brain. To support these visual functions, retinas across species share a basic cellular composition, comprising a resident glial cell type known as Müller glia and several major retinal neurons, including photoreceptors, bipolar cells, horizontal cells, amacrine cells, and retinal ganglion cells [1]. At the same time, retinal structure and function have evolved to adapt to diverse environmental demands, resulting in significant interspecies variations [2]. For instance, teleost fishes possess remarkable retinal regenerative potential, whereas mammalian retinas have lost this ability [3]. Nocturnal rodents have rod-dominated retinas for heightened sensitivity in dim light, while diurnal primates have evolved cone-rich macular regions for sharp vision [4,5]. Comparative studies across species not only deepen our understanding of conserved mechanisms in human retinal development but also provide insights into the pathogenesis of retinal diseases, such as age-related macular degeneration [6]. Beyond cross-species studies using naïve retinas, human retinal organoids, which mimic retinal structure and function, are increasingly used to model human retinal development and disease [7,8].

Single-cell sequencing technologies have advanced biomedical research, including retinal studies, by enabling comprehensive molecular profiling and mechanistic insights at the single-cell level [9,10]. Among these technologies, scRNA-seq and scATAC-seq are widely used to characterize single-cell transcriptomes and chromatin accessibility, respectively [11]. More recently, spRNA-seq has been developed to measure transcriptomes by incorporating spatial information at the level of individual spots or cells [12]. These technologies have provided profound insights into retinal biology across multiple dimensions. In cross-species studies, scRNA-seq has revealed the conservation and evolution of retinal cell types across 17 species [13], while scATAC-seq has identified conserved cell type-specific regulatory elements across species [14]. In retinal development, scRNA-seq has uncovered distinct developmental states and programs, with scATAC-seq revealing chromatin accessibility patterns that control temporal patterning, and spRNA-seq providing insights into spatiotemporal dynamics of cellular composition and cell-cell communication [10–13]. In retinal regeneration research, comparative analyses of species with different regenerative abilities using scRNA-seq have uncovered distinct molecular responses to retinal injury, revealing mechanisms that suppress neurogenic competence in mammalian species [15]. For retinal vascular diseases, integrative analysis of scRNA-seq and scATAC-seq has been employed to investigate pathological neovascularization shared across diabetic retinopathy, retinopathy of prematurity, and age-related macular degeneration, revealing distinct cell populations, changed chromatin accessibility, and gene regulatory networks underlying vascular pathologies [16]. However, despite the wealth of available data and the insightful utility of these technologies, current datasets are scattered across different repositories with inconsistent processing standards, limiting cross-study comparisons and systematic analyses. Most critically, there remains a lack of comprehensive, integrated databases that consolidate retina single-cell and spatial omics data under varied biological conditions across species, hindering the full potential of comparative genomics for understanding retinal biology and disease.

To address this gap, we developed scRetinaDB, a comprehensive database of single-cell and spatial sequencing retinal data across multiple species. scRetinaDB integrates 453 scRNA-seq datasets comprising over 2.79 million cells from 17 species and 27 biological conditions, alongside 107 scATAC-seq and 18 spRNA-seq datasets from human and mouse retinas. For each species, we integrated scRNA-seq data to construct a unified retinal cell atlas. scRetinaDB provides four main modules for browsing species-specific sequencing datasets, searching cross-species omics profiles, and applying analysis tools including cell type annotation and cell similarity comparison, respectively. Overall, scRetinaDB serves as a resource to support vision research by enabling systematic analyses of retinal single-cell and spatial sequencing data across species.

## Materials and methods

### Data collection

We systematically searched for and collected retina single-cell and spatial omics datasets from public repositories including Gene Expression Omnibus (GEO, https://www.ncbi.nlm.nih.gov/geo/), Genome Sequence Archive (GSA, https://ngdc.cncb.ac.cn/gsa/), and ArrayExpress (https://www.ebi.ac.uk/biostudies/arrayexpress), using keywords such as “retina”, “single-cell”, “scRNA-seq”, “scATAC-seq”, and “spatial RNA-seq”. A complementary literature search in the PubMed (https://pubmed.ncbi.nlm.nih.gov/) and Google Scholar (https://scholar.google.com/) was also performed using the same keywords. By manually reviewing each publication, we compiled detailed information of each dataset, including data accession, PubMed ID (PMID), sequencing platform, biological conditions, sample number, and cell counts. In total, the database includes three sequencing modalities (scRNA-seq, scATAC-seq, and spRNA-seq) collected from 17 different species. The data cover a wide range of biological conditions, including retinas from development, regeneration, and disease. Raw or processed sequencing data were downloaded from each original publication. To ensure data reliability and consistency, we performed unified data processing and quality control for each dataset.

### Single-cell RNA-seq data processing

For scRNA-seq data, we performed quality control directly on the gene expression matrices or raw count data. Raw sequencing data in SRA format were converted to FASTQ format using sratoolkit (v2.10.7), followed by alignment and quantification through the Cell Ranger (v7.1.0) pipeline. Taking human retinal data as an example, we collected data from 13 studies and performed quality control following these criteria: (1) each gene had to be detected in at least 10 cells; (2) only cells with 300–6500 total genes and 300–10,000 total reads were retained; (3) cells with mitochondrial gene expression exceeding 60% were removed; (4) doublets were identified and removed using DoubletFinder (v2.0.4); and (5) Mitochondrial genes were excluded from downstream analysis to avoid technical confounders and high variability that could interfere with accurate cell type annotation. The quality control criteria for other species are listed in Table S1.

### Single-cell ATAC-seq data processing

For scATAC-seq data, we preprocessed fragments in BAM format, which contained the genomic coordinates of ATAC-seq fragments associated with each cell. These fragments were used to generate feature matrices by quantifying the chromatin accessibility across various genomic regions, such as promoters, gene bodies, and distal regulatory regions. Using the Signac (v1.14.0) package [17], we created cell-by-peak count matrices, in which peaks were associated with the nearest genes based on genomic distance. Then, we followed the quality control criteria reported in the original studies (Table S2). For studies that did not report criteria for quality control, we implemented the standards recommended in the Signac documentation (https://stuartlab.org/signac/). Quality control typically involved the assessment of multiple metrics, including the number of accessible chromatin regions per cell, the count of sequencing fragments per cell, the percentage of sequencing reads within accessible chromatin regions, the proportion of reads in the blacklist of genomic regions (known to produce artifacts), the signal of nucleosome positioning, and the enrichment of accessibility signal around gene transcription start sites.

### Spatial RNA-seq data processing

For spRNA-seq data, gene expression matrices and corresponding H&E-stained images were processed using the Seurat package (v5.2.1) [18]. Spatial transcriptomic data were integrated into unified objects, and high-resolution images were aligned to gene expression spots. Low-quality spots with fewer than 10 counts were excluded from further analysis.

### Species-specific integration of single-cell RNA-seq datasets across biological conditions

We integrated scRNA-seq datasets for each species by correcting batch effects while preserving biological variation. Using human retinal datasets as an example, we first employed the merge function from the Seurat package to combine individual datasets. Data were then normalized using NormalizeData and subsequently scaled using ScaleData. Highly variable genes were identified using FindVariableFeatures (n = 2,000) and used for principal component analysis. Batch effects were corrected with Harmony (v1.2.3), incorporating three covariates with optimized lambda parameters (platform: 0.5; study: 1; sample: 2.5). Subsequent analyses were based on the first 24 Harmony-corrected dimensions, which were used for dimensionality reduction via uniform manifold approximation and projection (UMAP) using the RunUMAP function. Clustering was performed using FindClusters with a resolution of 0.1 for initial clustering, followed by re-clustering of major cell types at higher resolutions. Integration parameters for other species are included in Table S3, with adjustments to Harmony dimensions and clustering resolution.

### Cell type annotation

For each species in scRetinaDB, we included cell type markers curated from published literature. To annotate cell types of clusters using literature-curated markers in scRetinaDB, we developed a method which contains three major steps.

First, we calculated the power (*MP*_*c*, *t*, *g*_) of marker *g* from cell type *t* to distinguish cluster *c* and other clusters. The power was quantified based on AUC (area under the receiver operating characteristic curve of gene expression) calculated by the ROCR package in R platform. The relationship between MP and AUC is the following:

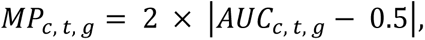

where *AUC*_*c*, *t*, *g*_ represents the area under the ROC curve for marker *g* from cell type *t* in cluster *c*.

Second, we computed the joint power (*JP*_*c*, *t*_) which measures the power of cluster *c* when assigned to cell type *t*. The formula was calculated as follows:

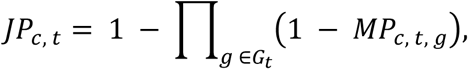

where *G*_*t*_ denotes the set of literature-curated markers from cell type *t*.

Finally, the cell identity of each cluster was determined based on the joint power *JP*_*c*, *t*_. The higher joint power indicates greater confidence that a cluster belongs to a given cell type. The cluster was annotated as the cell type with the highest joint power.

### Identification of marker genes in each cell type

For each species, cluster-specific marker genes were identified using the FindAllMarkers function in Seurat with the Wilcoxon rank-sum test. To ensure the comparability of marker gene profiles across datasets, the analysis was restricted to genes that were common across all datasets of a given species. Only positively enriched markers were considered, applying thresholds of at least 25% expression within a cluster and log2 fold-change > 0.5. We applied the Bonferroni correction to account for multiple hypothesis testing, and genes with an adjusted p-value < 0.01 were retained as significant markers of cell types.

### Cell-cell communication analysis

Cell-cell communication was analyzed for human and mouse retinal cell atlases, leveraging their well-established ligand-receptor databases. We employed the LIANA package (v1.0), which integrates multiple cell-cell interaction resources from CellPhoneDB, CellChat, and Connectome [19]. We analyzed cell-cell interactions separately across different studies and experimental conditions. In non-embryonic samples, we excluded progenitor cells (neurogenic progenitors, retinal progenitor cells, and proliferating retinal progenitor cells) to focus on mature cell populations. We identified significant interactions, defined as those with an aggregate rank ≤ 0.01 using the liana_wrap function and aggregated results across methods with liana_aggregate. The interactions were visualized through an interaction frequency heatmap, which displays the number of interactions between each pair of cell types.

### Analysis of cell type-specific regulatory activities

Gene regulatory networks were inferred only for human and mouse retinal datasets using pySCENIC (v0.12.1) [20]. The analysis consisted of three steps: (i) inference of gene regulatory networks using the GRNBoost2 algorithm, (ii) identification of transcription factor binding motifs through cisTarget analysis, and (iii) calculation of regulatory module activity scores using AUCell. Reference databases for motif analysis were downloaded from the cisTarget resources (https://resources.aertslab.org/cistarget/), including transcription factors and regulatory regions separately for mouse and human. Cell type-specific transcription factor activities were visualized using a heatmap.

### Database implementation

scRetinaDB was developed using a three-tier web application architecture consisting of the frontend, backend, and database layers. The frontend was built with Vue.js (v2.7.5), providing an interactive user interface through a single-page application (SPA). The backend employs Django (v5.0) running on Python 3.11 (Ubuntu 20.04.6) to handle business logic, data processing, and API services [21]. The deployment stack includes nginx (v1.22.1) as the reverse proxy server and uWSGI (v2.0.23) as the WSGI application server, enabling efficient request handling and communication between the web server and Django application. R (v4.3.3) is integrated into the Django backend for on-demand bioinformatics computations. The database uses MySQL (v8.1.0) as the relational database management system for storing expression data, metadata, analysis results, and reference information [22].

## Results

### Overview and statistics of the scRetinaDB

scRetinaDB comprises over 3.6 million retinal cells across 51 studies, including 453 scRNA-seq datasets, 107 scATAC-seq datasets, and 18 spRNA-seq datasets (Figure 1A, Table S4). The database spans 17 species, 8 sequencing platforms, and 43 biological conditions encompassing normal development, disease states, and regeneration.

**Figure 1.**
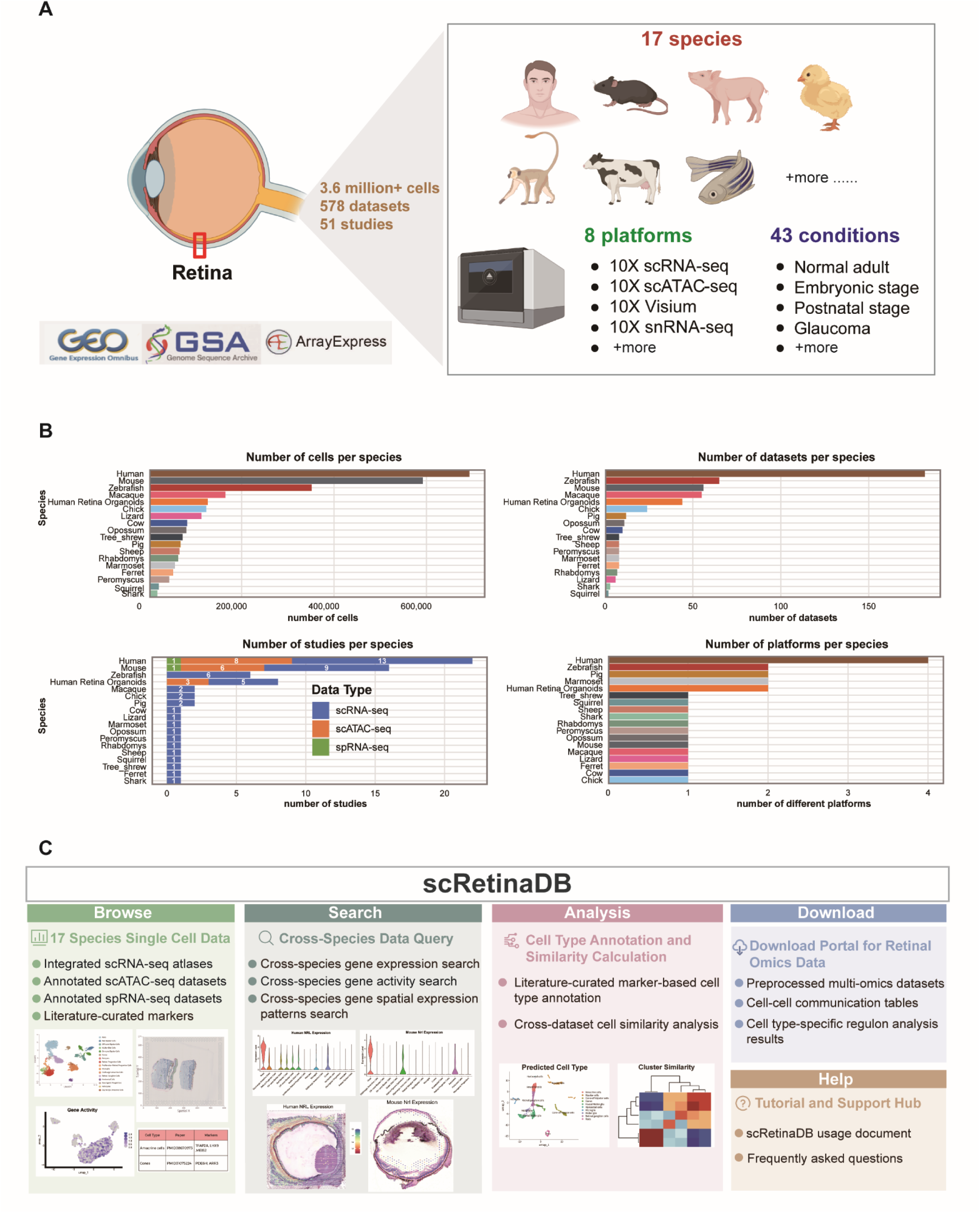
Overview of scRetinaDB database and user interface. Comprehensive collection of single-cell and spatial sequencing datasets covering 17 species, 8 platforms, and 27 biological conditions with 578 datasets from 51 studies totaling 3.6 million cells. **B.** Database statistics showing cell counts, dataset distribution, study numbers, and platform types across different species. **C.** scRetinaDB user interface providing browse, search, analysis, download, and help for integrated retinal datasets.

In scRetinaDB, scRNA-seq data cover 17 species, including human (native retinas and retinal organoids), macaque, marmoset, tree shrew, mouse, cow, pig, sheep, ferret, opossum, squirrel, peromyscus, rhabdomys, zebrafish, shark, chick, and lizard. scATAC-seq and spRNA-seq data are currently available for human (both native retinas and retinal organoids) and mouse. Species coverage varies by sequencing technology. Among all species, human, mouse, and zebrafish are the most extensively represented in terms of both cell counts and dataset numbers (Figure 1B).

To ensure data consistency across studies, we implemented standardized processing pipelines separately for the scRNA-seq, scATAC-seq, and spRNA-seq data (Figure S1). Each dataset undergoes rigorous quality control, normalization, and dimensionality reduction, followed by clustering and cell type annotation. Depending on data modality, downstream analyses include marker gene identification and cell-cell communication analysis for scRNA-seq data, accessible chromatin region and peak analysis for scATAC-seq data, and spatial domain analysis for spRNA-seq data. For paired scRNA-seq and scATAC-seq datasets, multi-omics integration analysis was performed to construct regulon networks.

The interface of the scRetinaDB website includes four major modules (Figure 1C). First, the Browse module allows users to explore integrated scRNA-seq atlases for each of 17 species, along with annotated scATAC-seq datasets and spatial RNA-seq datasets, including literature-curated cell type marker libraries. Second, the Search module enables cross-species queries of different omics data. Third, the Analysis module offers tools for marker-based cell type annotation and cell similarity calculation. Lastly, the Download module provides user-friendly access to preprocessed omics datasets and results from downstream analyses. scRetinaDB also provided a manual to help users navigate these resources.

### Species-specific retinal cell atlas derived from integrative scRNA-seq analyses

To characterize retinal cellular diversity, we constructed a cell atlas for each species through integrating scRNA-seq data across all biological conditions. We applied the Harmony algorithm to correct the batch effects arising from studies, sequencing platforms, and sample sources. Using human naïve retinal data, we assessed the effectiveness of the integrative analysis through UMAP visualization. Before integration, cells from different studies exhibited obviously separated clustering distributions, indicating strong batch effects across studies, while after integration, cells from different studies were well-aligned with the same cell types clustered in the same populations (Figure 2A). The similar distribution of normal retinal cells from different studies also indicated that the batch effects were removed after integration (Figure S2A). The integrated human retinal atlas revealed 16 distinct cell populations (Figure 2B), which showed specific expression of canonical retinal markers (Figure 2C). The integrative analysis revealed all major retinal cell types, including distinct subtypes of bipolar cells and amacrine cells. The cell atlas of naïve retinas included non-neuronal cell types, such as Müller glial cells, astrocytes, microglia, and pericytes. We also identified several progenitor cells during retinal development, such as retinal progenitor cells, proliferating retinal progenitor cells, and neurogenic progenitors.

**Figure 2.**
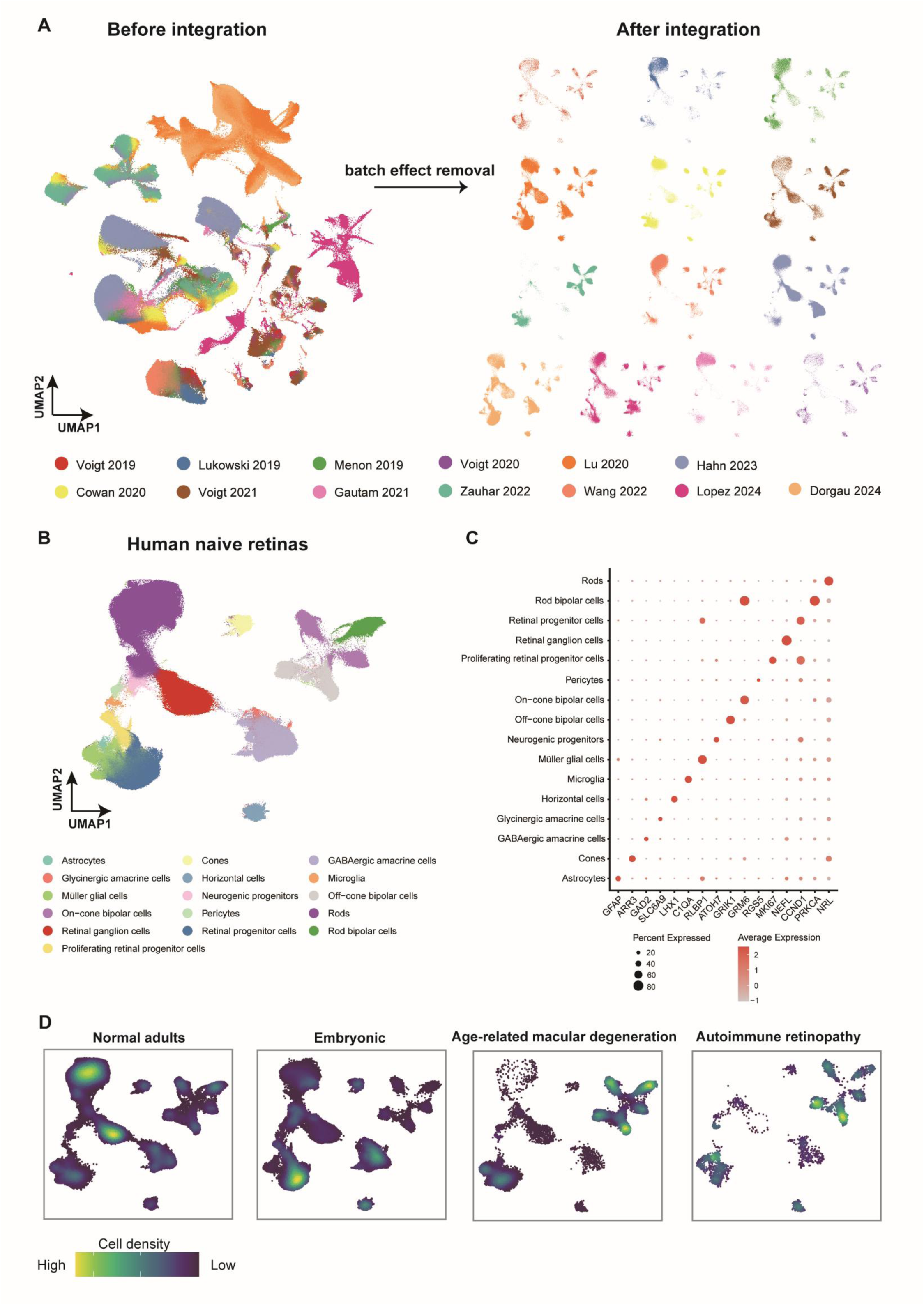
Human retinal cell atlas from integrating scRNA-seq datasets. **A.** UMAP visualization of naïve retinal samples before and after integration demonstrating effective batch effect removal across different studies. **B.** Integrated atlas identifying major retinal cell types and subtypes in human naïve retinas. **C.** Dot plot showing expression levels and percentages of canonical markers across retinal cell types. **D.** Density plots depicting cell distribution across normal adults, embryonic stage, age-related macular degeneration, and autoimmune retinopathy conditions.

We also examined the cell distribution across retinal development and disease in human retinal cell atlas (Figure 2D). Normal adult retinas contained all major cell types, with rods and retinal ganglion cells being particularly abundant (Figure 2D, S2B). Embryonic and early postnatal retinas contained substantial populations of retinal progenitor cells and proliferating retinal progenitor cells. Both AMD and autoimmune retinopathy (AIR) samples showed marked rod depletion, with AIR samples additionally exhibiting enrichment of microglia and reactive astrocytes as revealed by Ro/e analysis (Figure S2C). These cellular compositional changes in retinal degeneration diseases were consistent with known pathological features of photoreceptor degeneration [23,24]. The results demonstrated that our integration analysis for the species-specific retinal cell atlas removed technical batch effects while preserving biological variations across different physiological and pathological states.

Beyond the human naïve retinal cell atlas, we applied the same integration strategy to construct retinal cell atlas for each of all 17 species represented in scRetinaDB (Figure S3, S4). These retinal cell atlases provided a resource for cross-species comparisons, enabling identification of both conserved and species-specific programs in retinal organization and function.

### Analysis of fetal retinal development using scATAC-seq and spRNA-seq data

To illustrate scATAC-seq and spRNA-seq datasets in scRetinaDB, we performed omics analysis for human retinal development. By Integrating single-nucleus RNA sequencing (snRNA-seq) and single-nucleus ATAC sequencing (snATAC-seq) datasets publicly available from human fetal retinas [25], we dissected the regulatory mechanisms during retinal development from both transcriptomic and chromatin accessibility perspectives. UMAP analysis of multi-omics data revealed nine major retinal cell types consistently identified across both single-cell transcriptomes and chromatin accessibility profiles (Figure 3A). Analysis of cell type-specific regulon activities through integrating snRNA-seq and snATAC-seq datasets revealed the dynamic activation of transcriptional regulatory modules during fetal retinal development (Figure 3B). The *RAX* regulon exhibited activity in both progenitor cells and multiple differentiated cell types, reflecting its broad regulatory role in retinal development and maintenance. The *PAX6* regulon showed specific activation in amacrine cells, the *CRX* regulon displayed high activity in cone cells and bipolar cells, the *ISL1* regulon demonstrated strong activity in retinal ganglion cells, and the *LHX1* regulon was specifically activated in horizontal cells. These regulon activities illustrate the transcriptional regulatory basis underlying the transition from multipotent states to functionally specialized cells during fetal retinal development.

**Figure 3.**
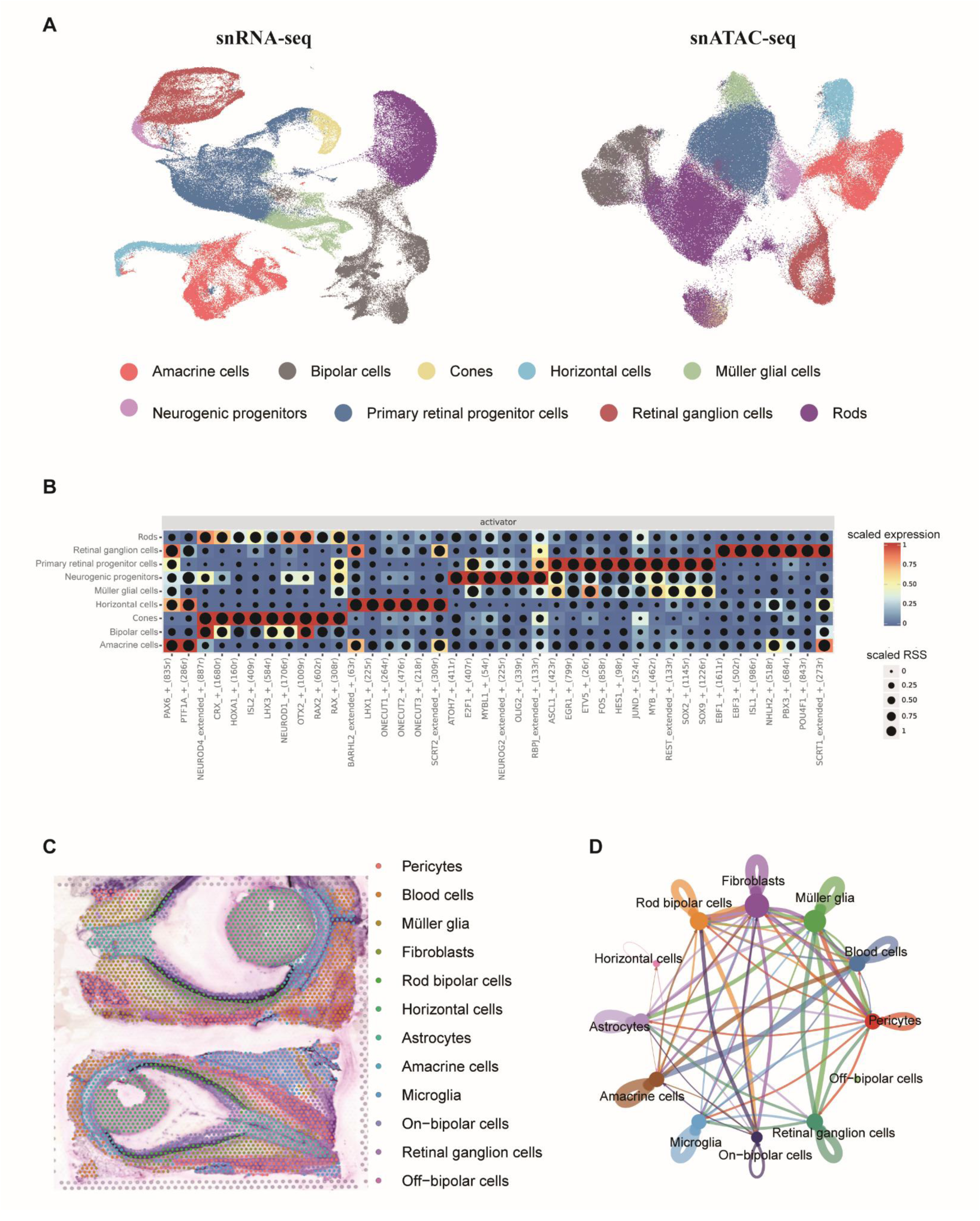
Single-cell multi-omics and spatial RNA-seq data analysis of human naïve retinas. **A.** Single-cell multi-omics atlas showing UMAP visualization of matched single-nucleus RNA sequencing (snRNA-seq) and single-nucleus ATAC sequencing (snATAC-seq) data from human retinal development (GSE268630). **B.** Cell type-specific regulon activity analysis based on integrated snRNA-seq and snATAC-seq data showing regulatory activities of transcription factors in different retinal cell populations. RSS, regulon specificity score. **C.** Annotation of the spatial RNA-seq dataset (HRA006282) from human retinas at 14 postconceptional weeks showing cell type distribution across tissue architecture. **D.** Spatial cell-cell communications of retinal cell types from HRA006282.

Spatial RNA-seq analysis of a representative fetal retina (HRA006282) revealed that by week 14 of gestation, retinal cells had begun to form a preliminary spatial distribution reflecting their eventual functional organization (Figure 3C). Spatially-informed cell-cell interaction analysis unveiled molecular communication networks between neighboring cell types during retinal development (Figure 3D). These local signaling exchanges may contribute to cell fate specification and tissue morphogenesis.

### Browse module for exploring single-cell and spatial sequencing data for the retina from each species

In scRetinaDB, we provide a Browse module to enable user-friendly access to single-cell and spatial sequencing data from each of the 17 species. The species information cards display comprehensive dataset statistics, including the number of cell types, total cell numbers, and available datasets, along with direct access to multi-omics resources. Using human retina as an example, the information card shows coverage of both native retinas and retinal organoids, providing access to the scRNA-seq cell atlas, scATAC-seq data, spRNA-seq data, and literature-curated markers (Figure 4A).

**Figure 4.**
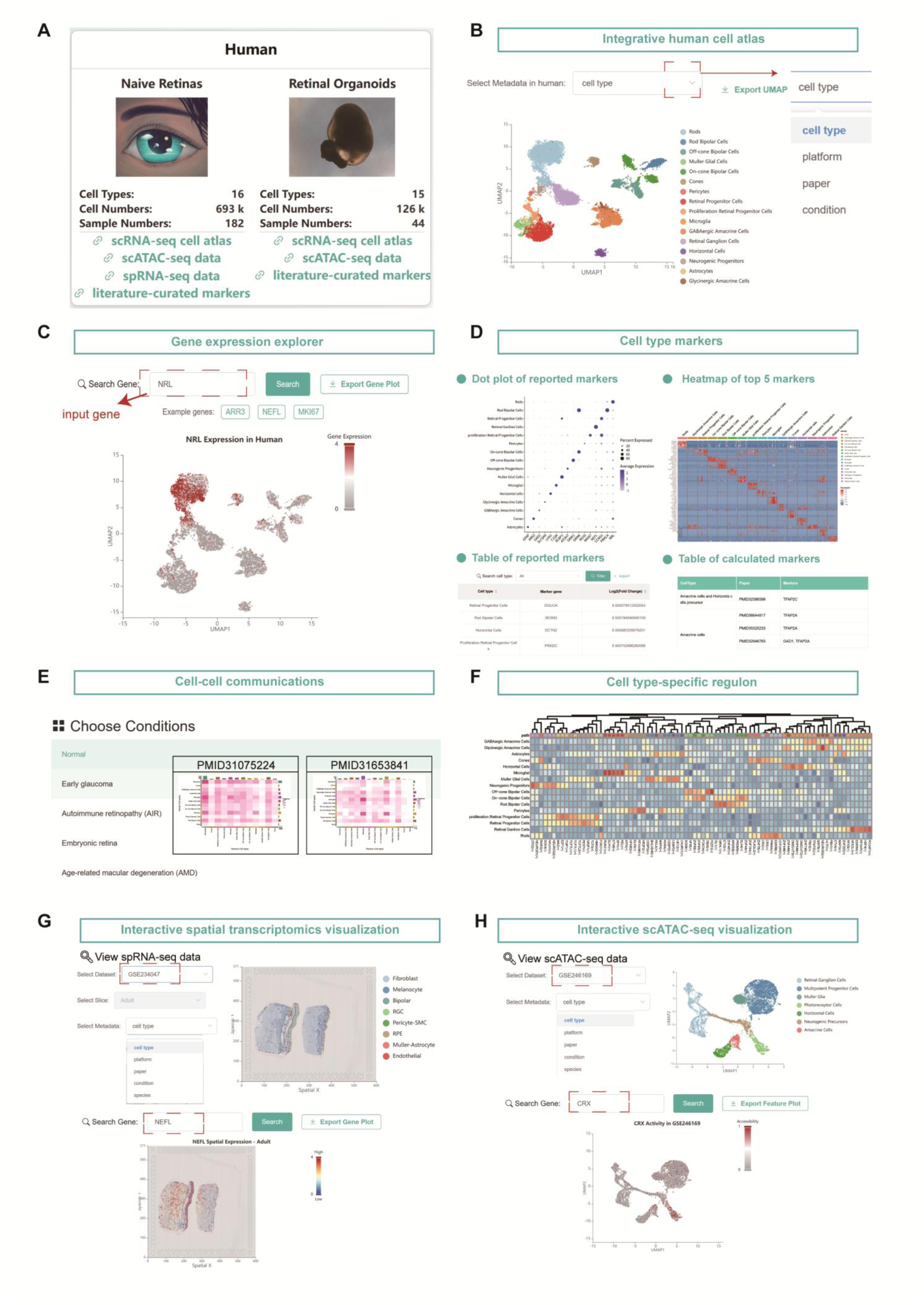
scRetinaDB browse interface using human retinas as an example. **A.** Human retina information cards showing dataset statistics and access links for naïve retinas and retinal organoids. **B.** Interactive UMAP visualization of integrated human scRNA-seq cell atlas with 16 cell types and metadata filtering options. **C.** Gene expression search interface showing *NRL* expression across the atlas. **D.** Cell type marker section including computationally identified markers with dot plots and heatmaps, and reported markers from literature. **E.** Cell-cell communication section showing intercellular interaction frequency heatmaps across different datasets and biological conditions. **F.** Cell type-specific regulon section showing transcriptional regulatory factors across cell types. **G.** Interactive spatial RNA-seq dataset with gene expression visualization (*NEFL* example). **H.** scATAC-seq dataset for chromatin accessibility analysis (*CRX* example).

Interactive UMAP visualization presents the integrated human scRNA-seq cell atlas generated through integrating all biological conditions from human naïve retinas (Figure 4B). Users can dynamically filter metadata to explore cellular populations according to cell type, sequencing platform, source publication, and experimental conditions, enabling comprehensive navigation of 16 distinct cell types identified in human retinas.

In the scRNA-seq cell atlas, we further provide multiple sections to show gene expression, marker genes, cell-cell communications, and cell type-specific regulons. The section for gene expression search allows users to query any gene across the integrated atlas (Figure 4C). Results are visualized as UMAP plots with expression intensity overlays, with *NRL* shown as an example. The marker gene section showcases computational identification of cell type-specific markers derived from the integrated cell atlas (Figure 4D). Visualization includes dot plots showing marker expression specificity across cell types and heatmaps displaying the top five markers for each cell type. Additionally, users can search, sort, and download marker genes through detailed marker gene tables that provide comprehensive information for each retinal cell type. Cell-cell communication section reveals intercellular interaction networks across different biological conditions using LIANA-based analysis (Figure 4E). This module displays condition-specific interaction frequency in normal, early glaucoma, autoimmune retinopathy (AIR), embryonic development, and AMD condition. The regulon section displays comprehensive heatmaps showing cell type-specific transcriptional regulatory factors and modules across human retinal cell populations (Figure 4F).

Beyond scRNA-seq data, scRetinaDB also provides access to spatial transcriptomics and scATAC-seq data with species and dataset selection capabilities. Spatial transcriptomics exploration offers interactive atlas browsing where users can search for specific genes (*NEFL* shown as an example) and visualize spatial expression patterns across retinal tissue architecture by integrating histological images with spatial transcriptome data (Figure 4G). Meanwhile, scATAC-seq data functionality allows users to query regulatory activities across cell types based on chromatin accessibility (*CRX* shown as an example) (Figure 4H).

### Analysis module for cell type annotation and cell similarity analysis

To facilitate cell type annotation using markers in scRetinaDB, we developed a computational tool represented in the Analysis module that supports cell type annotation for all 17 species (Figure 5A). Users can upload Seurat object files and select corresponding species through a user-friendly input interface. The prediction workflow utilizes literature-curated markers from scRetinaDB as references, implementing a three-step annotation process including assessment of marker specificity, modeling of combinatorial powers, and hierarchical annotation (Figure 5A). Higher power scores indicate a greater likelihood that a cluster belongs to a given cell type. Results are presented through both summary tables displaying predicted cell types with corresponding combinatorial powers for each cluster and interactive UMAP plots visualizing the distribution of annotated cell types for uploaded datasets.

**Figure 5.**
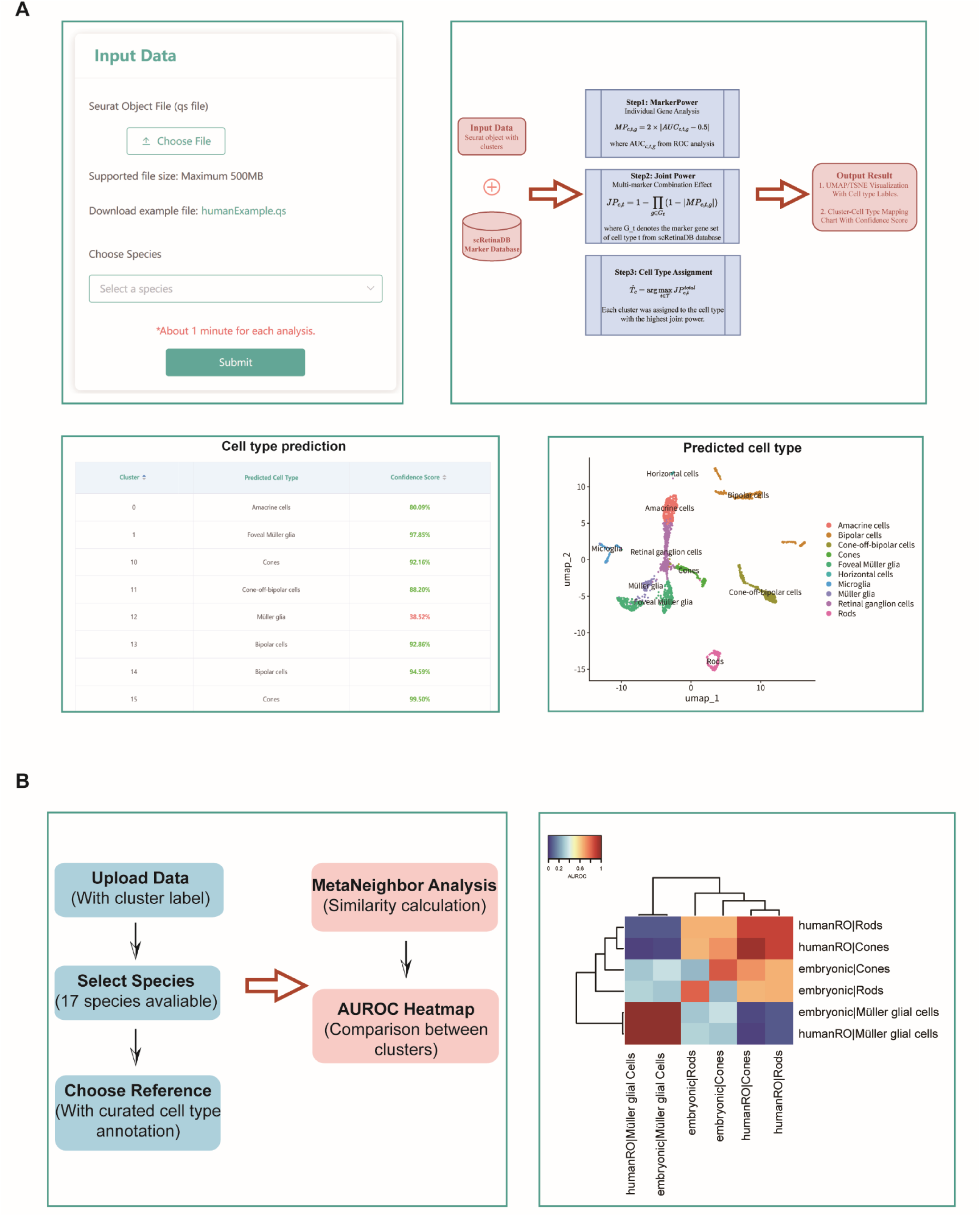
scRetinaDB analysis module for cell type annotation and cell similarity analysis. **A.** Cell type annotation tool interface and workflow. Input panel for uploading Seurat object files and species selection (left upper), three-step annotation algorithm workflow (right upper), prediction results showing cell types and confidence scores (left lower), and UMAP visualization of annotated cell types (right lower). **B.** MetaNeighbor similarity analysis workflow (left) and AUROC heatmap showing similarity scores between cell types from human retinal organoids and embryonic retinas (right).

The analysis module also incorporates MetaNeighbor for cell similarity calculation (Figure 5B). MetaNeighbor employs neighbor voting principles based on the concept that identical cell types should exhibit similar expression profiles, where predicted label distributions should approximate true label distributions for closely related cell types [26]. Users can upload datasets with cluster information, select target species, and choose reference datasets from scRetinaDB for similarity analysis. Results are presented as AUROC score heatmaps where deeper red coloring indicates higher similarity between cell types (Figure 5B). As demonstrated in the example comparing human retinal organoids with human embryonic retinal data, cones, rods, and Müller glia from organoid datasets show high similarity scores with corresponding embryonic cell types.

### Search module for cross-species comparison of retinal sequencing data

scRetinaDB offers a Search module to compare cross-species retinal sequencing data, enabling users to investigate molecular conservation and divergence among retinas from different species. The search interface supports multiple data types with species-specific coverage. For scRNA-seq data, users can search for gene expression across integrated atlases spanning all 17 species in the database. For scATAC-seq data, users can search for gene regulatory activities calculated from chromatin accessibility fragments, with support for human naïve retinas, human retinal organoids, and mouse retinas. Users can select specific datasets within each supported species for targeted comparisons (Figure 6A). The system automatically identifies homologous genes across target species and retrieves corresponding gene expression and regulatory activities. Results are presented through both UMAP and violin plot visualizations. UMAP plots display the integrated atlases in two panels, where cells are clustered based on transcriptional similarity and colored by cell type, with expression or activity intensity overlaid. Violin plots quantitatively represent distribution patterns across retinal cell types. As exemplified by the NRL search, the gene shows specific expression in rods in both human and mouse retinas, demonstrating comparative analysis capability across species.

**Figure 6.**
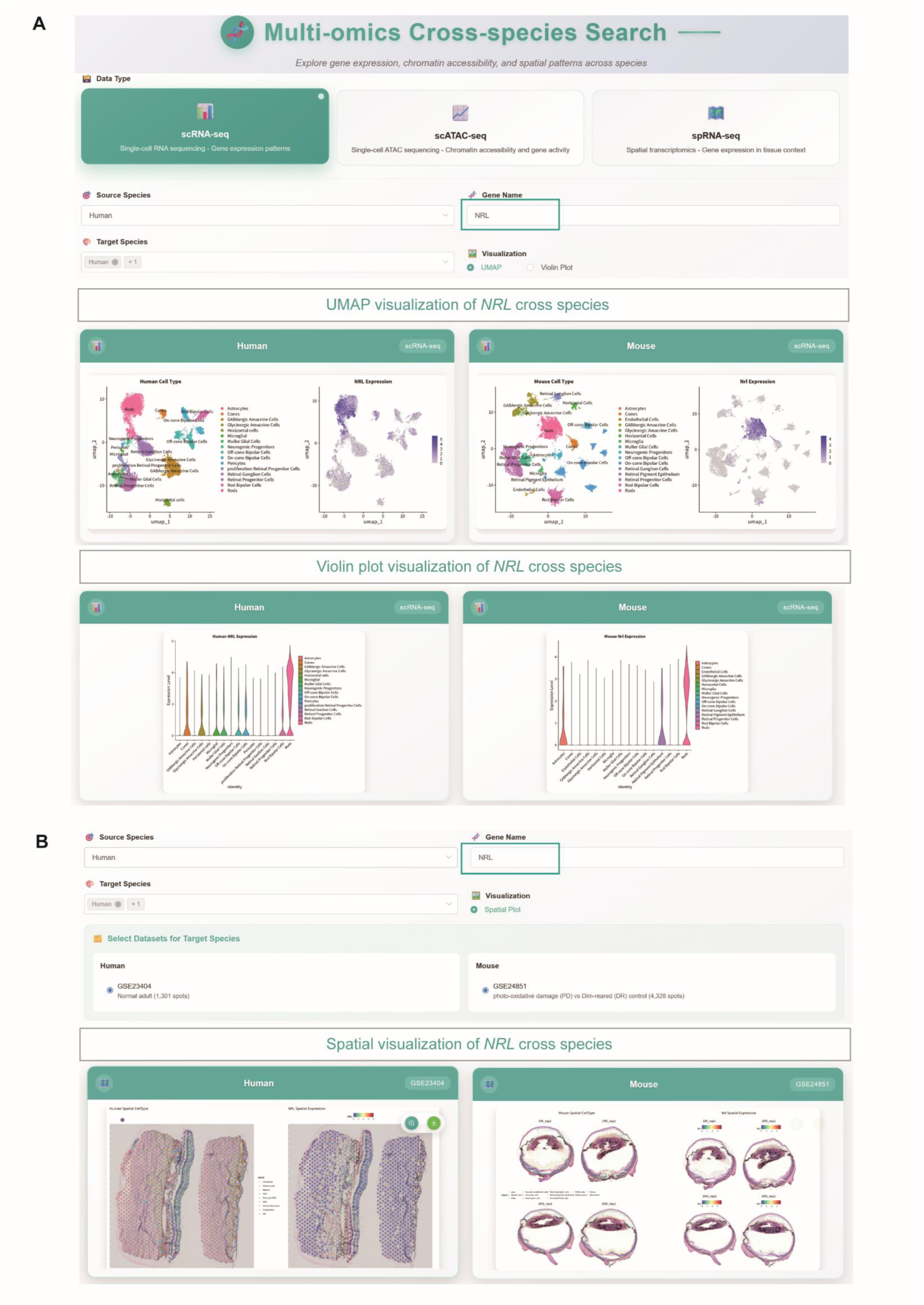
scRetinaDB cross-species search interface for data exploration. **A.** Gene expression or gene activity search interface with UMAP and violin plot visualizations for comparative analysis across 17 species. **B.** Spatial expression search tool for cross-species comparison of gene expression patterns in retinal tissue sections.

For spatial transcriptomics comparison, users can perform cross-species spatial expression analysis with dataset selection capabilities for human and mouse retinas (Figure 6B). As shown with NRL spatial expression, users can explore both spatial distributions of cell types and spatial gene expression patterns across retinal tissue sections, enabling comparative analysis of spatial expression domains and tissue organizations between species.

### Download module for the preprocessed datasets

To access and download high-quality retinal single-cell and spatial sequencing data, scRetinaDB provides a user-friendly Download module. The download interface is organized into scRNA-seq data, scATAC-seq data, spRNA-seq data, cell-cell communication table, and cell type regulon table.

The Download module uses a filtering system to identify subsets of datasets by setting publication year, sequencing platform, species, and biological condition (Figure 7A). Search results are displayed in a table including accession number, platforms, species, published year, PMID, condition, sample number, and cell number. Each dataset provides direct access to preprocessed data files, which are available in compressed formats compatible with standard analysis frameworks such as Python/Scanpy.

**Figure 7.**
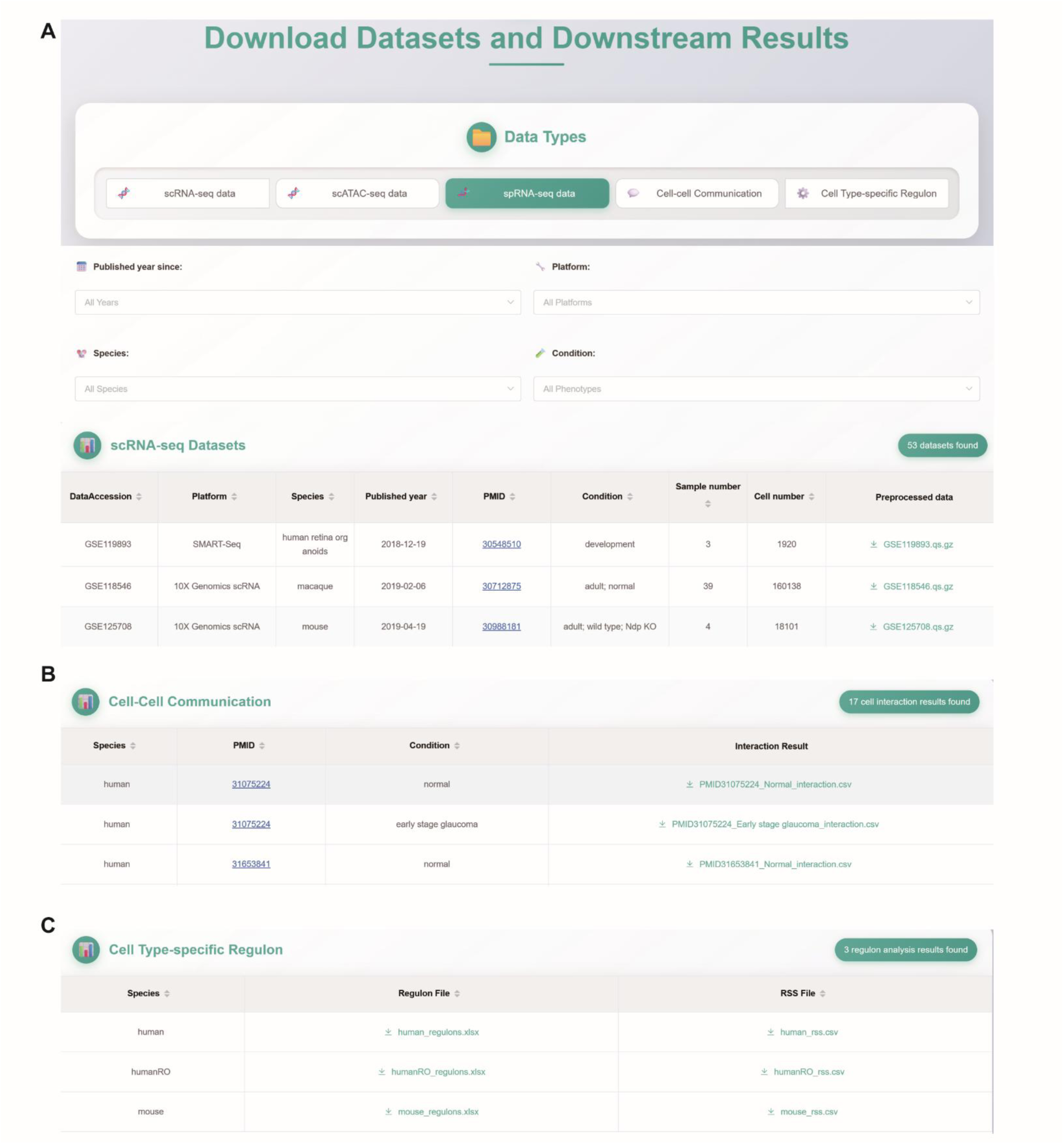
scRetinaDB download portal for datasets and analysis results. Interface showing downloadable datasets and analysis results separately from scRNA-seq datasets with metadata and preprocessed files (**A**), cell-cell communication analysis (**B**), and cell type regulon analysis (**C**) across different species and conditions.

Beyond raw datasets, scRetinaDB also provides downloadable analysis results corresponding to the analyses displayed in the Browse module. Cell-cell communication tables are organized by species, with corresponding PMID references and condition-specific interaction files available for download (Figure 7B). Cell type regulon tables provide species-specific regulons and RSS (regulon specificity score) for transcriptional regulatory network analysis (Figure 7C). These downloadable analysis results correspond to the computational outputs presented in the cell-cell interaction heatmaps and regulon visualizations within the Browse module.

## Discussion

In this study, we presented scRetinaDB, a comprehensive database that aggregates single-cell and spatial omics data alongside literature-curated marker genes from retinal tissues across multiple species and biological conditions. The database covers 17 vertebrate species and several biological conditions, including retinal development, regeneration, and disease. For scRNA-seq data, encompassing over 2.79 million cells from 453 datasets of 34 studies, we established retinal cell atlas spanning different biological conditions for each species. Additionally, scRetinaDB incorporates scATAC-seq datasets with over 820,000 cells and spRNA-seq datasets with more than 30,000 spots from human and mouse retinas. Users can browse and download profiles from each sequencing modality, as well as downstream analyses such as cell type-specific regulons and cell-cell communications. The Search module enables cross-species comparisons, while the Analysis module provides tools for cell type annotation and single-cell similarity calculation.

Advances in sequencing technologies, particularly single-cell and spatial sequencing technologies, have generated extensive retina datasets. Previously, only two retina-related omics databases were available: RETINA, which compiles RNA-seq data from human retinas (https://retina.tigem.it/), and iSyTE, which provides microarray expression data from mouse and human retinas (https://research.bioinformatics.udel.edu/iSyTE/) [27,28]. However, these existing databases are primarily based on traditional bulk sequencing or microarray technologies, lacking single-cell and spatial sequencing data. They typically cover only two species, and are unable to support comprehensive cross-species comparative analyses. Most importantly, they lack systematic integration of multiple sequencing modalities across diverse biological conditions such as development, disease, and regeneration. To our knowledge, scRetinaDB represents the first comprehensive database dedicated to retinal single-cell and spatial sequencing data across species and biological conditions. Beyond hosting scRNA-seq, scATAC-seq and spRNA-seq datasets, scRetinaDB also offers species-specific retinal cell atlases generated by integrating scRNA-seq data across diverse sequencing platforms and biological conditions. These atlases provide single-cell-level insights into the conserved and divergent mechanisms underlying retinal development, regeneration, and disease.

Using scRetinaDB, users could perform diverse omics studies across multiple levels. For example, the database facilitates omics studies of retinal development, regeneration, and disease, especially from a cross-species perspective. For cross-species analysis, several tools, such as CACIMAR and CAME, are available [29,30]. Integration of multi-omics data in scRetinaDB will consolidate insights from individual omics studies into a broader view of retinal biology. In addition, the datasets in scRetinaDB could serve as valuable training resources, especially for deep learning applications [31].

Given the rapid development of sequencing technologies and the growing retinal omics data, we are committed to continuously update scRetinaDB. In the future, scRetinaDB will incorporate additional retinal data modalities, such as single-cell DNA methylome sequencing [32]. Further analysis tools for retinal omics data will be developed and integrated into scRetinaDB. Altogether, scRetinaDB provides the retinal single-cell and spatial sequencing data and analysis tools that could advance vision research, particularly cross-species studies of retinal development, regeneration, and disease.

## Supporting information

Quality control criteria for scRNA-seq data across different species.

Study-specific quality control criteria for processing scATAC-seq data.

Harmony integration and clustering parameters in different species.

Summary of all datasets included in scRetinaDB.

## Declarations

### Ethics approval and consent to participate

Not applicable.

### Consent for publication

Not applicable.

### Availability of data and materials

scRetinaDB is a database for single-cell and spatial omics data from retinal tissues across 17 species (https://casapp.dnayun.com/scretina/). All public scRNA-seq, scATAC-seq, and spRNA-seq datasets used during the current study are available in Table S4. All preprocessed datasets can be downloaded from the Download page in the scRetinaDB.

### Competing interests

The authors declare that they have no competing interests.

### Funding

This study was supported by the National Natural Science Foundation of China [No. T2222003, 32570985, 32170849], the Ministry of Science and Technology of China [No. 2025YFC3409301, 2022YFA1105400], the Guangdong Province Science and Technology Program [No. 2023B1212060050, 2020B1212060052], and the National R&D Infrastructure and Facility Development Program of China [DKA2017-12-02-22].

### Authors’ contributions

JW, XX, FJ, and MT conceived and designed the project. MT, KY, and JC collected the data. MT, FJ, and ZC designed and constructed the database and toolkit. XX, AFS, JC and JLW tested the database website. MT, XX, and AFS analyzed and interpreted the data. MT prepared the manuscript. JW, XX, FJ, AFS, and JLW reviewed and edited the manuscript. JW acquired funding support. All authors read and approved the final manuscript.

## Acknowledgements

We sincerely thank all contributing projects and studies for providing the valuable resources utilized in scRetinaDB. Species images used in the web interface were obtained from Pixabay (www.pixabay.com). The human retina illustration was obtained from Unsplash (www.unsplash.com).

---

**Figure S1.**
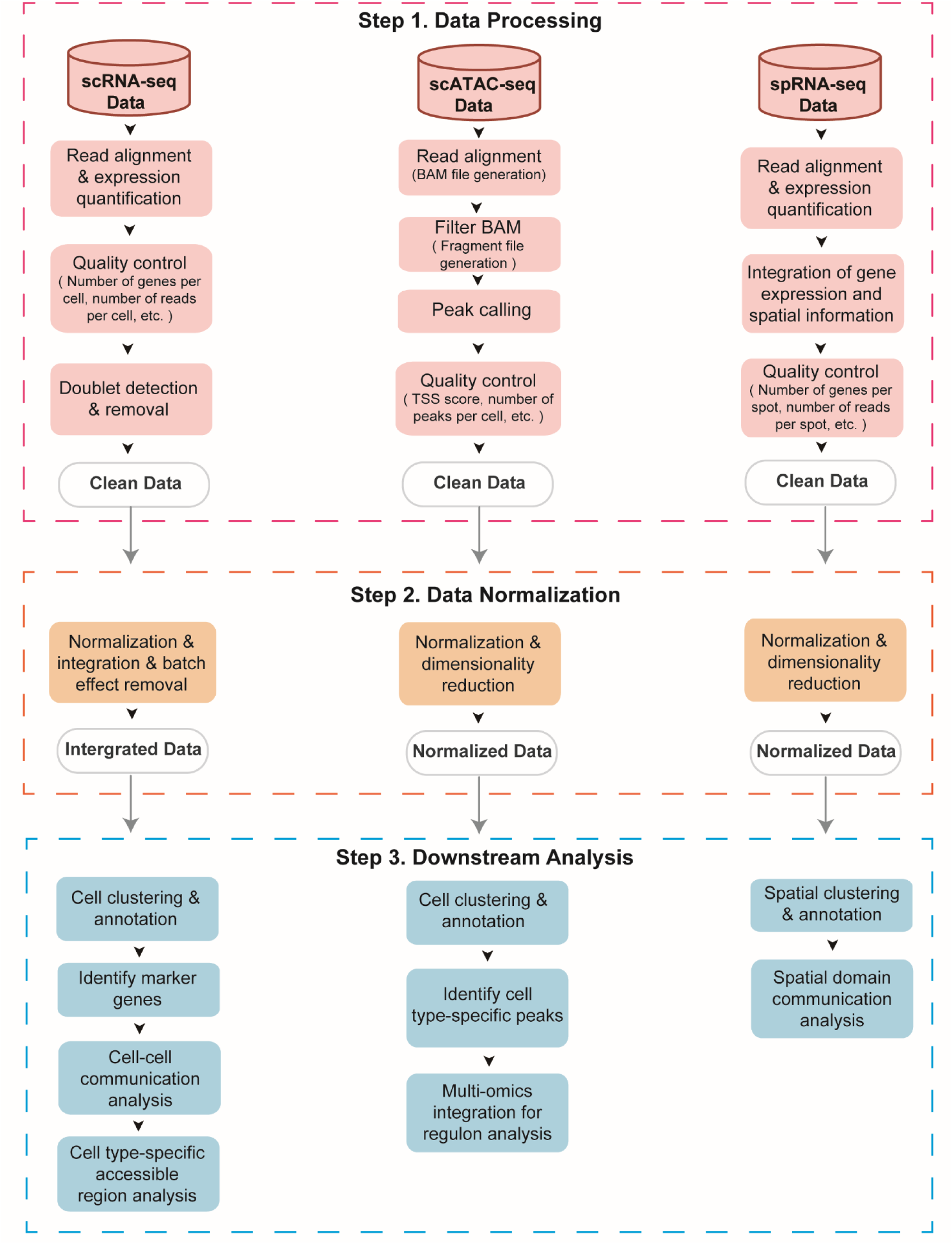
Standardized data processing and integration workflow for scRNA-seq, scATAC-seq, and spRNA-seq datasets. The workflow includes three main steps: data processing with quality control and alignment, data normalization and standardization, and downstream analysis including clustering, annotation, cell-cell communication and cell type-specific regulon analysis.

**Figure S2.**
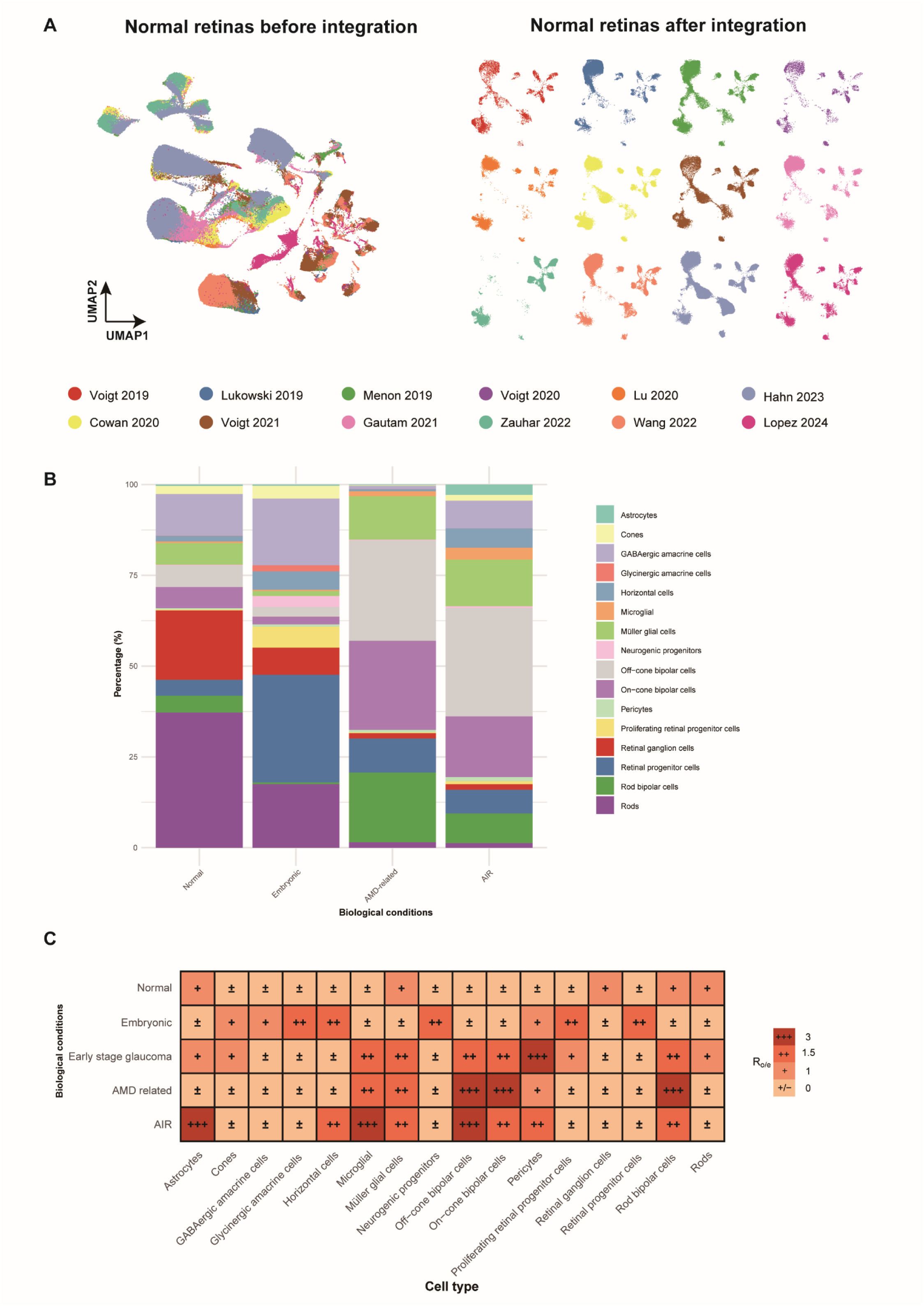
Cell type distribution and enrichment analysis across different retinal conditions. **A.** Clustering of human naïve retinas from normal adult before and after integration using Harmony software. **B.** Stacked bar chart showing the percentage of major retinal cell types in normal, embryonic, early glaucoma, AMD, and autoimmune retinopathy (AIR) samples. **C.** Heatmap showing Ro/e (ratio of observed to expected) analysis across different retinal conditions and cell types, with color intensity and asterisks indicating significance levels.

**Figure S3.**
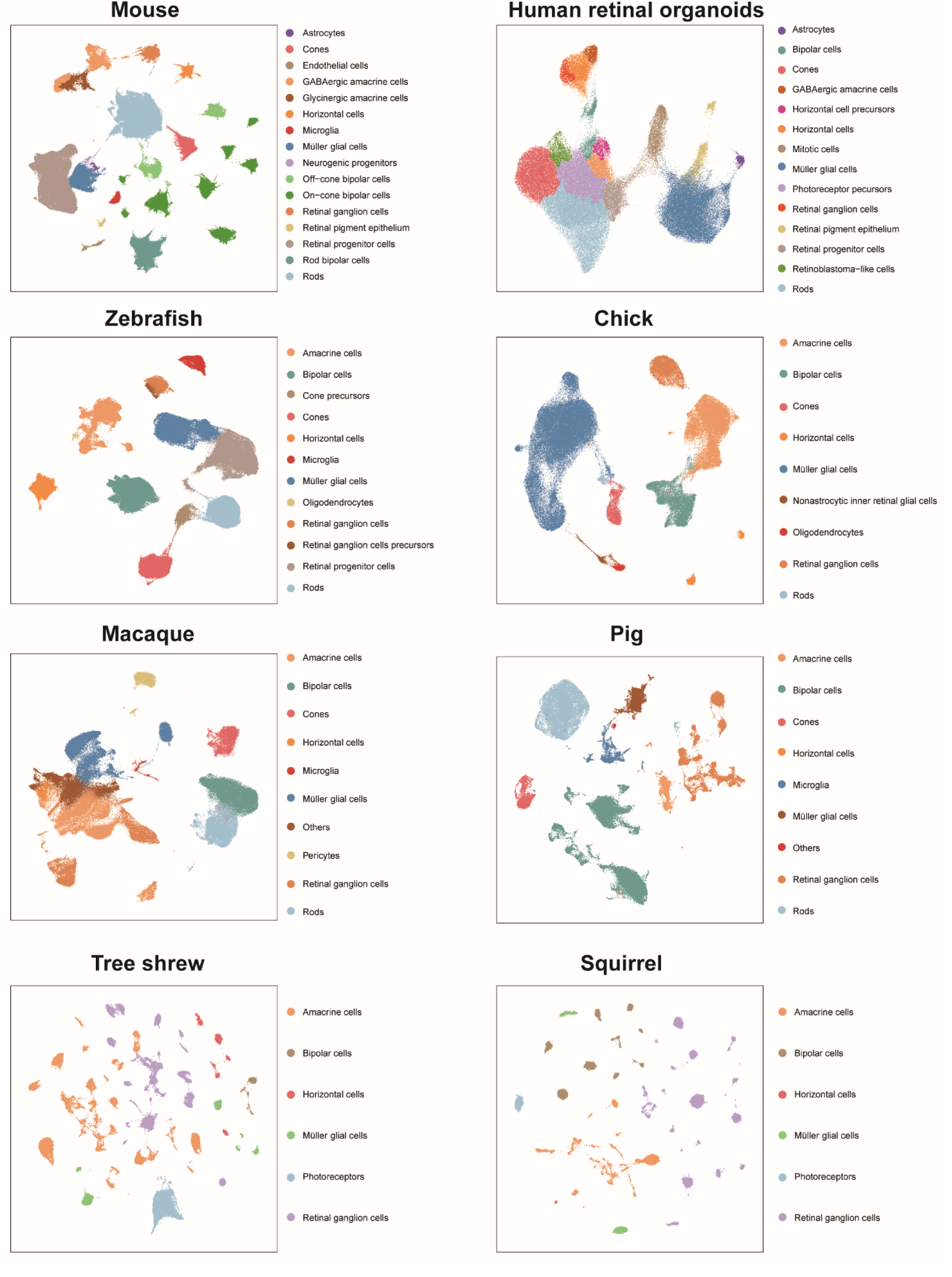
Retinal cell atlases from eight species. For each species, scRNA-seq datasets from all biological conditions were integrated.

**Figure S4.**
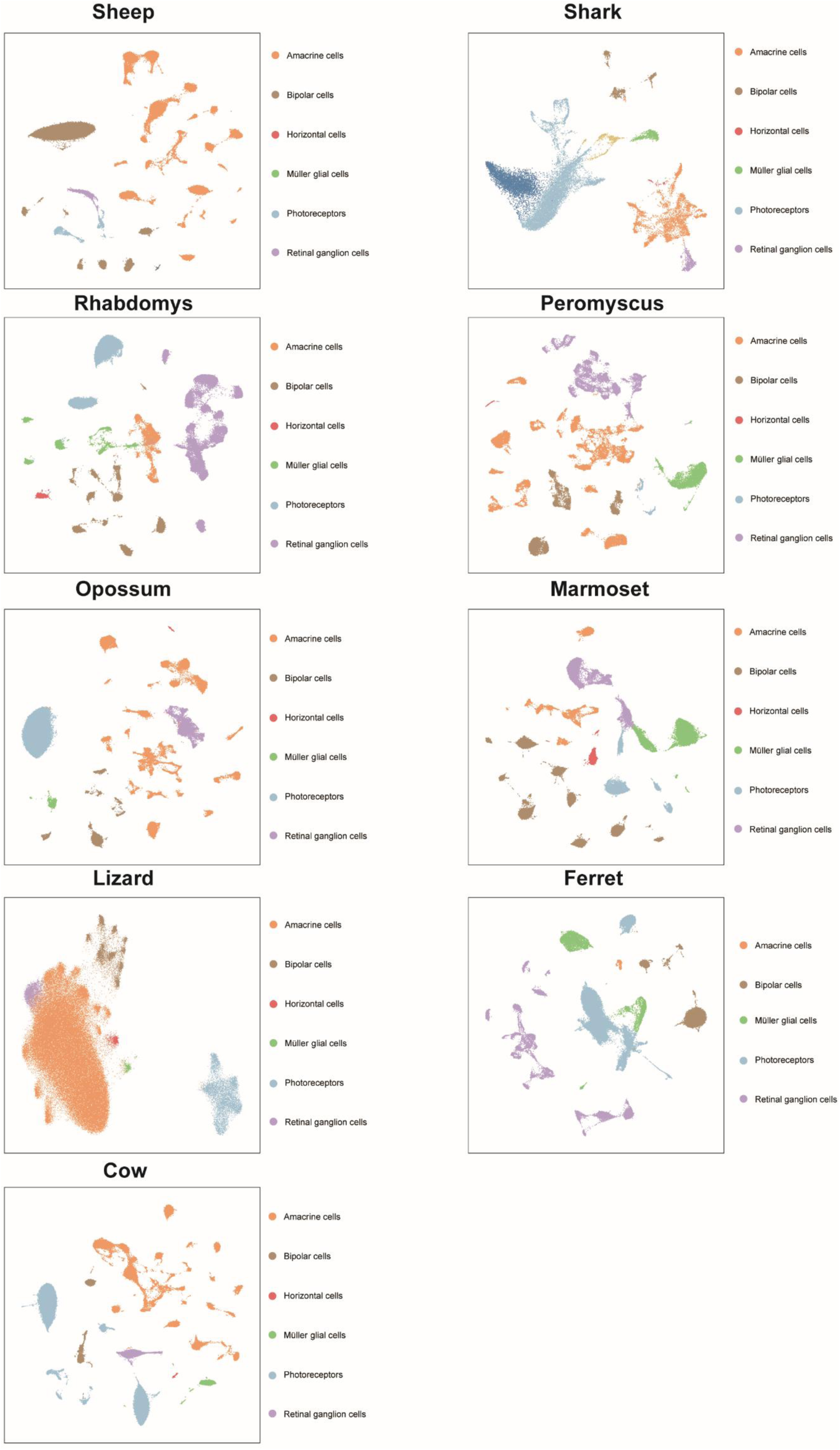
Retinal cell atlases from nine species. For each species, scRNA-seq datasets from all biological conditions were integrated.

## Table legends

**Table S1: Quality control criteria for scRNA-seq data across different species.**

**Table S2: Study-specific quality control criteria for processing scATAC-seq data.**

**Table S3: Harmony integration and clustering parameters in different species.**

**Table S4: Summary of all datasets included in scRetinaDB.**

## Notes

https://casapp.dnayun.com/scretina/

### Competing Interest Statement

The authors have declared no competing interest.

